# Complex Interactions Between Genome Components of Two ToLCNDV Isolates from India and Spain Determine Effective Infection in Tomato

**DOI:** 10.64898/2025.12.11.691045

**Authors:** Verónica Pérez-Rubio, Isabel Fortes, Beatriz Romero-Rodríguez, Laura Arribas-Hernández, Enrique Moriones, Araceli G. Castillo

## Abstract

Tomato leaf curl New Delhi virus (ToLCNDV) is a bipartite begomovirus whose infectivity and host range depend on the coordinated functions of its DNA-A and DNA-B components. Here, we investigated the replication, movement, and systemic infection capacity of ToLCNDV isolates from India (IN) and Spain (ES) and their pseudo-recombinants in tomato and *Nicotiana benthamiana*. In tomato, the IN isolate (A-IN/B-IN) systemically infected all plants, whereas the ES isolate (A-ES/B-ES) failed to do so. Pseudo-recombinant analyses revealed that DNA-B from the IN isolate complemented A-ES in trans, enabling systemic spread, while B-ES accumulated poorly and supported systemic infection only inefficiently. Although both A components replicated locally in tomato, DNA-A from the IN isolate was unable or only marginally able to infect systemically in the absence of DNA-B, demonstrating an essential contribution of DNA-B to long-distance movement. In *N. benthamiana*, both isolates established systemic infections even as monopartite viruses, though with substantially reduced viral titers and attenuated symptoms, indicating that host factors can partially compensate for the absence of DNA-B. Using a DsRed-tagged ΔCP mutant of ToLCNDV-ES, we further show that coat protein is required for systemic movement in *N. benthamiana*, despite the presence of DNA-B-encoded movement functions. Collectively, these results uncover striking host- and isolate-dependent differences in component compatibility and demonstrate that systemic infection by ToLCNDV-IN DNA-A in tomato requires DNA-B, while ToLCNDV-ES additionally depends on coat protein for efficient movement in *N. benthamiana*.

## Introduction

Begomoviruses (family *Geminiviridae*, genus *Begomovirus*) are plant-infecting DNA viruses with single-stranded circular genomes encapsidated in twinned icosahedral particles. They are exclusively transmitted by *Bemisia tabaci* whiteflies and cause major crop losses worldwide, particularly in tropical, subtropical and temperate regions (Kumar 2019; Rojas et al. 2018). Begomoviruses have a phylogeographic distribution in which most viruses classified as Old World (OW) species possess a monopartite genome (approximately 2.8 kb), whereas New World (NW) species mostly possess a bipartite genome (two components with a total size of approximately 5.2 kb) (Zerbini et al. 2017). Monopartite begomoviruses are phloem limited, with those from the OW usually associated with small circular ssDNA satellites. In contrast, bipartite begomoviruses have two 2.6 kb DNA components (DNA-A and DNA-B) which are separately encapsidated and both required for successful infection (Zhou 2013). The bipartite genome organization, thought to have arisen early in begomovirus diversification, is associated with distinct evolutionary dynamics between the two components. The evolutionary origin of DNA-B is still under investigation, but a widely discussed “capture hypothesis” proposes that it arose from a satellite molecule that was domesticated by a monopartite ancestor. Over time, DNA-B became indispensable for bipartite begomoviruses, supplying essential intra- and intercellular movement functions. This scenario aligns with the marked genomic and functional asymmetries observed between DNA-A and DNA-B: while DNA-A encodes proteins required for replication, transcriptional regulation, defense suppression and encapsidation, such as the Replication-associated protein (Rep), Transcriptional activator protein (TrAP), Replication enhancer protein (REn), C4 and Coat protein (CP), the DNA-B encodes two movement-related proteins, the Nuclear shuttle protein (NSP) and Movement protein (MP). The single genomic component of monopartite geminiviruses is homologous to the DNA-A component of bipartite viruses and several of its proteins facilitate intercellular movement and systemic infection. Moreover, OW monopartite begomoviruses typically encode an additional gene, V2, which functions as both a movement protein and an RNA silencing suppressor (Fondong 2013; Kamal et al. 2024) and recent studies have identified novel ORFs, such as C5, C6 and V3, potentially involved in host manipulation and defense suppression(Gong et al. 2021; Wang et al. 2022; Zhao et al. 2023). Due to the limited capacity of the geminivirus capsid and their compact genome organization with little scope for additional genes, the evolutionary path for the monopartite progenitors of modern bipartite begomoviruses involved genome expansion through acquisition of an independently encapsidated DNA, either a second genomic component (DNA-B) or a satellite DNA.

Although the OW begomoviruses are primarily monopartite, there are species that possess a bipartite genome, including *Tomato leaf curl New Delhi virus* (ToLCNDV), *Pepper yellow leaf curl Indonesia virus*, and *Tomato yellow leaf curl Kanchanaburi virus* (Bagewadi and Naidu 2016; Sakata et al. 2008; Zaidi et al. 2017). Among these, ToLCNDV stands out for its exceptionally broad host range and extensive geographic distribution (Moriones et al. 2017; Zaidi et al. 2017). Typical symptoms of the Tomato leaf curl New Delhi disease include upward leaf curling, vein thickening, internode shortening, stunting, and yield loss when infection occurs early (Sahu et al. 2010). First identified in India on tomato in 1995 (Padidam et al. 1995), ToLCNDV now infects at least 43 dicot species spanning *Solanaceae*, *Cucurbitaceae*, *Malvaceae*, *Euphorbiaceae*, and *Fabaceae*, as well as multiple weed hosts (Juárez et al. 2019; Srivastava et al. 2016; Venkataravanappa et al. 2018, 2019; Zaidi et al. 2016). It has spread from the Indian subcontinent to the Middle East, North Africa, and southern Europe, where it causes severe epidemics in both open-field and greenhouse crops (Fortes et al. 2016; Juárez et al. 2014, 2019; Mnari-Hattab et al. 2015; Panno et al. 2019; Sifres Cuerda et al. 2018; Zaidi et al. 2017). Phylogenetic analyses suggest that Mediterranean populations likely arose from a single founder introduction from Asia(Zaidi et al. 2017). (Fortes et al. 2016) demonstrated that the majority of ToLCNDV isolates from Asian regions cluster into a single phylogenetic group of closely related DNA-A sequences, sharing ≥94% nucleotide identity, despite the broad diversity of host species, geographic origins, and sampling years. In contrast, ToLCNDV isolates emerging in the western Mediterranean Basin (ToLCNDV-ES strain) were shown to form a distinct phylogenetic group with DNA-A sequences sharing more than 99% identity among themselves, but less than 94% identity with other known ToLCNDV DNA-A sequences.

Interestingly, although most ToLCNDV strains infect a range of Cucurbitaceae hosts, distinct host specificities are evident among isolates. Asian strains typically infect both Solanaceae (e.g., tomato, eggplant, chili) and Cucurbitaceae (e.g., zucchini, cucumber, melon), whereas Mediterranean isolates (e.g., from Spain and Italy) are primarily adapted to cucurbits and display poor or no infectivity in tomato under natural conditions. However, published reports on this matter are inconsistent. Several studies state that Mediterranean isolates fail to infect tomato or do so only at very low efficiency (Fortes et al. 2016; Vo et al. 2023), whereas others claim that isolates belonging to the same Mediterranean strain readily infect tomato (Janssen et al. 2022; Ruiz et al. 2015; Yamamoto et al. 2021).

Unlike most bipartite begomoviruses, ToLCNDV from the Asian isolates is considered a non-obligate bipartite virus, its DNA-A can establish infection without the cognate DNA-B in some hosts, and it can form pseudo-recombinants by associating with DNA-B components from other begomoviruses (Iqbal et al. 2025; Zaidi et al. 2016). Although satellites are not essential for symptom development, ToLCNDV can associate with diverse betasatellites (e.g., chilli leaf curl betasatellite, radish leaf curl betasatellite) that can intensify pathogenicity or partially compensate for DNA-B loss(Agnihotri et al. 2018; Akhter et al. 2014; Jyothsna et al. 2013; Shahid et al. 2021; Vignesh et al. 2023). This notable capacity for diverse viral interactions may play a key role in shaping its broad host range and wide geographical distribution(Iqbal et al. 2025; Zaidi et al. 2017). Genetic analyses involving targeted mutations of all ToLCNDV genes have been conducted to elucidate their specific roles in viral replication, movement, and pathogenicity (Padidam et al. 1995, 1996). While the CP of canonical New World (NW) bipartite begomoviruses has been shown to be dispensable for systemic movement (Brough et al. 1988; Gilbertson et al. 2003; Pooma et al. 1996; Sudarshana et al. 1998), mutations in the CP gene of ToLCNDV do not affect symptom development but severely reduce viral single-stranded DNA (ssDNA) accumulation in *Nicotiana benthamiana* and tomato (Padidam et al. 1995, 1996). These findings highlight functional differences between NW and Old World (OW) bipartite begomoviruses regarding the role of CP during infection.

The molecular determinants that enable ToLCNDV to infect tomato, as well as the factors underlying the inability of certain geographic strains to establish effective infections, remain poorly understood. Given ToLCNDV’s ability to associate with diverse satellites and other begomoviruses, its unusual non-obligate bipartite nature, and its variable host adaptation, identifying these determinants is critical for understanding host specificity, predicting virus emergence, and developing durable resistance in tomato. Notably, the vast majority of studies on ToLCNDV have focused on Asian isolates, whereas much less is known about the biology, host adaptation, and pathogenicity of isolates of the ToLCNDV-ES strain. In this study, we address this question by comparing two ToLCNDV isolates with differential abilities to infect tomato: one from India and one from Spain. We assessed their infectivity in tomato and *Nicotiana benthamiana* using pseudo-recombinants derived from both viruses. Our results highlight the critical role of genetic interactions between the two genomic components in establishing a successful infection. We found that the Spanish ToLCNDV isolate replicates efficiently in tomato but fails to establish systemic infection when paired with its cognate DNA-B, even though this DNA-B can provide, in trans, the movement proteins that enable the DNA-A from the Indian isolate to systemically infect tomato. Furthermore, by replacing the CP gene with DsRed without altering the overlapping AV2 gene, we generated a ΔCP clone for the Spanish isolate. Using this construct, we demonstrated that, at least in *Nicotiana benthamiana*, ToLCNDV-ES does not behave as a canonical bipartite begomovirus: the coat protein is essential for viral movement and systemic infection. Moreover, the movement proteins encoded by DNA-B appear to enhance, rather than replace, CP function.

## Material and methods

### Geminiviral infectious clones

Infectious clones of ToLCNDV-ES were generated from the DNA-A and DNA-B genomic components of the isolate ToLCNDV-[ES-Alm-661-Sq-13] (GenBank accession numbers KF749223 and KF749226, respectively), as previously described by (Fortes et al. 2016). For the ToLCNDV isolate from India, belonging to the ToLCNDV-India strain group (hereafter referred to as ToLCNDV-IN), the infectious tandem-repeat clones of DNA-A (GenBank HM007120) and DNA-B (GenBank U15017) were kindly provided by Prof. Supriya Chakraborty (Molecular Virology Laboratory, School of Life Sciences, Jawaharlal Nehru University, New Delhi, India).

Regarding the_infectious clone of the partial deletion of the coat protein (CP) gene (AV1) of the ToLCNDV-ES (DNA-A^ΔCP^) was constructed by inserting the *DsRed* fluorescent protein to generate a transcriptional fusion between the gene for the CP and *DsRed*. The starting material consisted of a dimeric DNA-A-ES clone previously inserted into the pBluescript SK + (pBSSK) vector (Stratagene, La Jolla, CA, USA)(Fortes et al. 2016). The monomeric DNA-A was excised with *BamH*I and recloned into pBSSK to generate pBSSK-DNA-A-ES vector. To remove the C-terminal region of the AV1 gene, a PCR amplification was performed over pBSSK-DNA-A-ES using Q5 high-fidelity DNA polymerase (New England Biolabs, Ipswich, MA, USA). The entire plasmid was amplified except for the 483 bp corresponding to the CP C-terminal region. The primers CP-Cter-*Hpa*IFw (5′-CCCCGTTAACGGCCTGTACTCATGCATCA-3′) and AV2-*Hpa*IRv (5′-CAGCGTTAACCCTTGGCACGTCAGGACT-3′) introduced *HpaI* restriction sites flanking the amplified fragment. In parallel, the *DsRed* coding sequence, was amplified from pBSSK-DsRed using the primers *Hpa*I-DsRedFw (5′-CTCTGTTAACATGGCCTCCTCCGAGAACG-3′) and *Hpa*I-DsRedRv (5′-GCCGGTTAACCTACAGGAACAGGTGGTGG-3′), which also incorporated *HpaI* restriction sites. The monomeric pBSSK-DNA-A-ES vector and the *DsRed* fragment were both digested with *Hpa*I (Takara, Shiga, Japan) and joined using T4 DNA ligase (New England Biolabs, Ipswich, MA, USA) to generate DNA-A^ΔCP^::*DsRed* construct.

To generate the infectious 1.2-mer construct, an additional 613 bp fragment containing the common region (CR) and part of the N-terminal region of the replication-associated protein (C1) was excised by digesting the dimeric clone with *BamH*I and *Xba*I and subsequently cloned into pBSSK to generate pBSSK-C1IRV2. Then, the previous DNA-A^ΔCP^::DsRed construct was digested with *BamH*I and the fragment containing the DNA-A^ΔCP^::DsRed sequence was ligated with the pBSSK-C1IRV2 vector also digested with *BamH*I, yielding the modified DNA-A clone (DNA-A^ΔCP^::DsRed 1.2 mer). Finally, the full-length DNA-A^ΔCP^::DsRed 1.2 mer was subcloned into the binary vector pCAMBIA0380 using PCR-amplified fragments generated with primers *Spe*I-1.2merFw (5′-TCGGACTAGTGGATCCCACATGTTTGTGGA-3′) and *Spe*I-1.2merRv (5′-CACAACTAGTAACGTCTCCGTCTTTGTCGA-3′), followed by *Spe*I digestion and ligation. The resulting construct was sequenced using Oxford Nanopore Technology (Plasmidsaurus. Cologne, Germany) to confirm the sequence integrity.

All the geminiviral infectious clone were introduced into *Agrobacterium tumefaciens* strain C58C1 using the freeze–thaw transformation method described by (Weigel and Glazebrook 2006).

### Plant and virus inoculation

*Solanum lycopersicum* (cv. Moneymaker) and *Nicotiana benthamiana* seeds were pre-germinated for 48 h at 25 °C in darkness and then transferred to growth chambers maintained at 25 °C and 60 % relative humidity, with a 16-h photoperiod under 120nmol·s⁻¹·m⁻² photosynthetically active radiation.

*Agrobacterium tumefaciens* cultures harboring each viral clone were grown in 15 mL of LB broth supplemented with kanamycin (50 µg/mL) and rifampicin (50 µg/mL) at 28 °C for 24 h with shaking. Bacterial cells were collected by centrifugation and resuspended in infiltration buffer (10 mM MgCl₂, 10 mM MES, 1 mM acetosyringone) to an optical density at 600 nm (OD₆₀₀) of 1.0. *A. tumefaciens* strains carrying the recombinant plasmids ToLCNDV-IN-A/ToLCNDV-IN-B, ToLCNDV-ES-A/ToLCNDV-ES-B, ToLCNDV-IN-A/ToLCNDV-ES-B, and ToLCNDV-IN-B/ToLCNDV-ES-A were mixed in a 1:1 ratio. The resulting cell suspensions were used in infectivity assays on three-week-old plants, either by stem puncture for systemic infection assays or by leaf agroinfiltration for local replication assays. Between 3 and 5 biological replicates were performed for each infection assay using from 6 to 8 plants per condition (see each figure legend for details).

### Sample collection and nucleic acid extraction

Agroinfiltrated leaves from tomato or N. benthamiana were collected at 5 days post-infiltration (dpi) to assess the replication ability of the infectious clones (local infection) . To evaluate the systemic infection, apical leaves were sampled at from 21 to 35 dpi depending on the assay (see each figure legend for details). Infected plants were shock-frozen in liquid nitrogen and ground to a fine powder using a mortar and pestle. Genomic DNA was extracted from either 60 mg (for PCR) or 1 g (for Southern blot) of leaf tissue. All reagent volumes described below correspond to the extraction of 60 mg of tissue. For the 1 g samples, the same protocol was followed but all volumes were scaled proportionally to the tissue mass. The extraction was carried out as follows: 400 µL of extraction buffer (100 mM Tris-HCl pH 8.0, 50 mM EDTA pH 8.0, 500 mM NaCl) were added to each sample and mixed thoroughly by vortexing. Subsequently, 80 µL of 10% SDS were added, and the mixture was incubated at 65°C for 10 min. After incubation, 160 µL of 5 M potassium acetate were added, and samples were placed on ice for 10 min. The lysates were centrifuged at 13,000 rpm for 20 min, and the supernatant was transferred to a new microcentrifuge tube. The aqueous phase was extracted with equal volumes of phenol and chloroform:isoamyl alcohol (24:1), mixed by vortexing, and centrifuged for 5 min at 13,000 rpm. The supernatant was transferred to a new tube and re-extracted with an equal volume of chloroform:isoamyl alcohol (24:1). DNA was precipitated with 700 µL of isopropanol and centrifuged at 13,000 rpm for 10 min. The resulting DNA pellet was washed with 70% ethanol, air-dried, and resuspended in nuclease-free water. DNA quality and concentration were determined by spectrophotometry and agarose gel electrophoresis.

### Detection of viral DNA by hybridization

Agroinoculated plants were analyzed by squash blot hybridization as previously described by (Fortes et al. 2016).

The probes used for the Southern blot hybridization, were based on ToLCNDV clones and the digoxigenin (DIG)-labelled DNA probes were prepared by mixed polymerase chain reaction (PCR) as described by (Navas-Castillo et al. 1999). For the DNA-A component the primer pair MA1788 (5′-CGTGTCGTTTCGATCTGGTGTC-3′) and MA1789 (5′- GTTTGTGGATCTAAACTTGGTGAG-3′), designed for the intergenic region (IR) of the DNA-A component of ToLCNDV-ES, which also hybridizes with the IR from the Indian isolates, was used (Fortes et al. 2016). A second probe was generated using the primer pair 2478 (5′-GCTTTTCCTTCTCCTTATTCCACTC-3′) and 2479 (5′-CCGTAAAGTCCATTTGTTTGAACAAC-3′), targeting the NSP gene of the DNA-B component of ToLCNDV-ES, which also hybridizes with the NSP gene from the Indian isolates, was used (Fortes et al. 2016).

Total DNA (1 µg) was electrophoresed on 0.8% agarose gels containing 0.5 µg/ml ethidium bromide in 0,5× TBE buffer for approximately 2 h, until the tracking dye reached the end of the gel. Gels were visualized under UV illumination to verify DNA integrity. For Southern blot transfer, gels were treated for 15 min in 0.25 M HCl with gentle shaking to depurinate high-molecular-weight DNA, followed by three rinses in distilled water. DNA was then denatured for 30 min in denaturation solution (1.5 M NaCl, 0.5 M NaOH) and neutralized for 30 min in neutralization solution (1.5 M NaCl, 0.5 M Tris-HCl, pH 7.2). After a brief rinse in distilled water, gels were equilibrated in 20× SSC buffer. After capillary transfer After capillary transfer of DNA onto a nylon membrane by upward capillarity in 20× SSC, following the standard procedure described by (Mamiatis et al. 1985), membranes were prehybridized for 2 h at 65 °C in prehybridization buffer (5× SSC, 1% blocking reagent, 0.1% N-lauroylsarcosine, 0.02% SDS, and 0.1 M NaCl) with rotation. Before hybridization, the probe was denatured by boiling the tube at 100 °C for 10 min. The denatured probe was then added to the prehybridized membranes, and hybridization was carried out overnight at 65 °C. Following hybridization, membranes were washed twice in 2× SSC containing 0.1% SDS at room temperature for 5 min, followed by three washes in 0.5× SSC containing 0.1% SDS at 65 °C for 10 min each. Membranes were finally rinsed once in maleic acid buffer (0.1 M maleic acid, 0.15 M NaCl, pH 7.5). For immunological detection, membranes were incubated for 30 min in blocking buffer (1% blocking reagent in maleic acid buffer) and then for 1 h with anti-digoxigenin-alkaline phosphatase conjugate (Roche) diluted 1:10,000 in the same buffer. After two washes (15 min each) in washing buffer (maleic acid buffer containing 0.3% Tween-20), membranes were equilibrated for 5 min in detection buffer (0.1 M Tris-HCl, 0.1 M NaCl, 0.05 M MgCl₂, pH 9.5). Chemiluminescent signals were developed using CDP-Star (Roche) according to the manufacturer’s instructions and visualized with a ChemiDoc™ Imaging System (Bio-Rad).

### Detection of viral DNA by PCR and qPCR

Primers were designed to amplify each DNA component of the bipartite ToLCNDV genome. PCR reactions were performed using GoTaq® DNA Polymerase (Promega, Madison, WI, USA) following the manufacturer’s instructions. The primer sets used for amplification were 2682 (5′-AGACTACGATCAAAGAAACCCC-3′) and 2683 (5′-TTCTTCTGAATTTAAACTTCAGCCC-3′) for the B-IN component; 2478 (5′-GCTTTTCCTTCTCCTTATTCCACTC-3′) and 2479 (5′-CCGTAAAGTCCATTTGTTTGAACAAC-3′) for the BES component; 2583 (5′-CAGCCCCTATGGAGCTCGTGCAGT-3′) and 2584 (5′-GAAATCCTGGGGAGATCCTGTA-3′) for the AIN component; 2585 (5′-ACAGCCCCTATGGAACCCGTG-3′) and 2586 (5′-TCCCTTGATGCATACGTTCCT-3′) for the AES component; 2480 (5′-GGATCCATTATTGCACGAATTTCCG-3′) and 2481 (5′-CATAGGGGCTGTCGAAGTTGAGCC-3′) for ΔCP; and 25SrRNA ITS-F (5′-ATAACCGCATCAGGTCTCCA-3′) and 25SrRNA ITS-R (5′-CCGAAGTTACGGATCCATTT-3′) as the endogenous control (Internal Transcribed Spacer, ITS).

Quantitative PCR (qPCR) reactions were performed using SsoFast™ EvaGreen® Supermix (Bio-Rad Laboratories, Hercules, CA, USA) according to the manufacturer’s instructions. Each reaction contained 10 ng of total DNA as template. The DNA-A from ToLCNDV-ES was amplified with the primer pair ToLCNDV-ES-Fw-qPCR (5′-CGTCCTACTATCGCCCTCTATGA-3′) and ToLCNDV-ES/IN-Rv-qPCR (5′-ATTTCATCCTTCGACAGAGTTCC-3′). These primers specifically amplify the DNA-A component of the ES strain, whereas the following forward primer ToLCNDV-IN-Fw-qPCR 5′-CGTTCTACTATCCCCCTCTATGA-3′), together with the same reverse primer, amplifies the corresponding region of the DNA-A component of the IN strain. Relative viral accumulation were performed by applying the 2^−ΔΔCt^ method.

For PCR and qPCR, the internal transcribed spacer (ITS) region of the 25S rRNA gene was used as an endogenous reference gene in *Nicotiana benthamiana* and *Solanum lycopersicum* samples.

### RCA and sequence analysis

Genomic DNA from tomato samples was extracted as described above. Rolling circle amplification (RCA) was carried out using the TempliPhi™ DNA Amplification Kit (Cytiva) according to the manufacturer’s instructions, with 1 µL of genomic DNA (40ng) as template in a final reaction volume of 10 µL. The reaction was incubated at 30 °C for 18 h to amplify circular viral genomes. RCA products were checked by restriction analysis but sent without digestion or additional purification for sequencing, using Oxford Nanopore Technology (Plasmidsaurus. Cologne, Germany).

### Confocal images

For images of *N. benthamiana*, leaf tissues were collected at 3 dpi from agroinfiltrated areas or 21 dpi for apical tissue. Confocal imaging was performed on a ZEISS LSM 880 confocal microscope equipped with a 10× objective. DsRed fluorescence was captured using sequential scanning with excitation at 561 nm and emission detected between 579–709 nm. Chloroplast autofluorescence was recorded using the same excitation wavelength (561 nm) and emission range.

## Results and Discussion

### The DNA-B component from ToLCNDV-IN provides, in trans, the functions required for the ToLCNDV-ES DNA-A component to achieve systemic infection in tomato

Tomato cv. *Moneymaker* plants were agroinfiltrated with infectious clones derived from DNA-A and DNA-B components of the Spanish (A-ES and B-ES) and Indian (A-IN and B-IN) isolates of ToLCNDV, as well as with the pseudo-recombinants A-IN/B-ES and A-ES/B-IN (Figure 1A).

**Figure 1.**
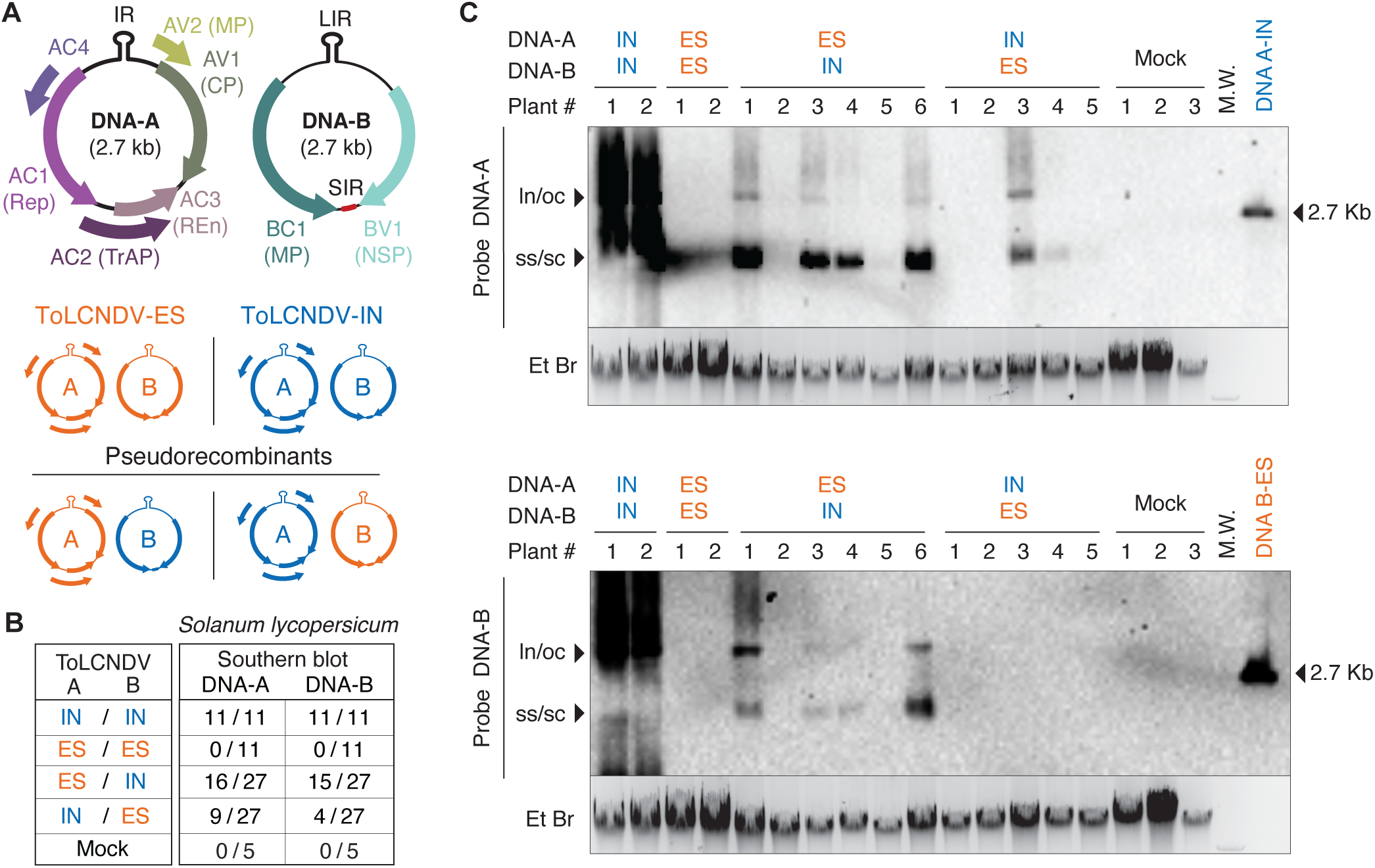
Detection of ToLCNDV IN and ES isolates and their pseudo-recombinants in tomato by Southern blot. (A) Genome organization of ToLCNDV. The ORFs (AV1, AV2, AC1, AC2, AC3, AC4, BV1 and BC1) and their protein products (CP, capsid protein; MP, movement protein; Rep, replication-associated protein; TrAP, transcriptional activator protein; REn, replication enhancer protein; C4, C4 protein; NSP, nuclear shuttle protein) are shown. IR, intergenic region; LIR, long intergenic region; SIR, short intergenic region; CR, common region. The hairpin which includes the origin of replication is indicated in the IR/LIR. Representation of the pseudo-recombinants between the IN (in blue) and the ES ToLCNDV isolates (orange) used in this work (below). (B) Summary of the viral detection in apical tomato leaves at 21 days post-infection (dpi) by Southern blot using DNA-A and DNA-B specific probes that recognize both the IN and ES ToLCNDV isolates. The table shows the number of plants testing positive for viral DNA relative to the total number of inoculated plants across three biological replicates. For each replicate, sample sizes ranged from 2 to 4 plants for the ToLCNDV-IN or ToLCNDV-ES isolates, 8 to 12 plants for each pseudo-recombinant, and 2 to 3 plants for the mock control (binary vector lacking viral sequences). (C) Southern blot detection of viral DNA in apical tomato leaves at 21 dpi. Approximately 1 µg of total genomic DNA per sample was separated on 0.8% agarose gels containing ethidium bromide, transferred to nylon membranes, and hybridized with DNA-A or DNA-B specific probes that detect both the IN and ES ToLCNDV isolates. The positions of the single-stranded (ss), supercoiled (sc), open circular (oc), and linear (ln) viral DNA forms, as well as the molecular-weight (M.W.) marker, are indicated. As an additional size reference, *Xba*I digested fragments from DNA-A-IN or DNA-B-ES infectious clones that released the corresponding monomeric viral genomic component (∼2.7 kb) were included on the right side of each gel. Samples shown correspond to one representative biological replicate.

Symptom development was monitored across five independent biological replicates (R1–R5). As previously reported (Fortes et al. 2016), all tomato plants inoculated with ToLCNDV-IN (A-IN/B-IN) developed severe symptoms, including stunting, leaf curling, crinkling, interveinal yellowing, and vein clearing (Figure S1). In contrast, plants inoculated with ToLCNDV-ES (A-ES/B-ES) remained asymptomatic. Among the pseudo-recombinants, A-ES/B-IN did not induce symptoms in any of the 58 assayed plants, while 8.6% (5/58) of the plants inoculated with A-IN/B-ES developed severe symptoms comparable to those observed with the IN isolate (Table S1, Figure S1).

To assess systemic infection and viral accumulation, apical leaves were collected, and viral DNA was analysed using squash blot hybridization from cross-sections of petioles (biological replicates R1 and R2) or Southern blotting (R3, R4 and R5). In plants infected with the IN isolate, viral DNA from both components was consistently detected in all samples (20/20 by squash blot; 11/11 by Southern blot), although DNA-B accumulated at lower levels (Figure 1C). By contrast, no viral DNA was detected in systemic tissues of plants inoculated with the ES isolate (0/20 by squash blot; 0/11 by Southern blot) (Figures 1B, 1C, and S2).

When the pseudo-recombinant A-ES/B-IN was inoculated, viral DNA was detected in most of the infected plants (17 of 20, analysed by squash blot, and 16 of 27 of those determined by Southern blot, all asymptomatic) (Figures 1B, 1C, and S2). In the case of the pseudo-recombinant A-IN/B-ES, viral DNA was detected in lower proportion, 25% of plants by squash blot (5/20, three symptomatic) and in 33% by Southern blot (9/27, two symptomatic) (Figures 1B, 1C, and S2). Importantly, symptom development correlated with the presence of the A component from the IN isolate, although not all A-IN/B-ES inoculated plants showed symptoms. Overall, regardless of the genomic combinations, the accumulation of DNA-B was consistently lower than that of DNA-A, or even undetectable in some plants infected with the pseudo-recombinants (Figures 1B and 1C).

Detection of the A-ES component in systemic leaves of A-ES/B-IN–inoculated plants (33/47; 70.2%), coupled with its absence in A-ES/B-ES–inoculated plants (0/31), indicates that A-ES can replicate in tomato but fails, or only poorly succeeds, in establishing a systemic infection with its cognate B-ES (Figures 1B, 1C, and S2).To confirm this hypothesis, we performed local infection assays and analysed the viral-agroinfiltrated leaves at 5 dpi. Southern blotting revealed replicative forms of A-ES regardless of the B component present in the leaf. Moreover, A-ES supported the replication of heterologous DNA-B, although less efficiently than A-IN (Figure 2). Significantly, B-ES was only weakly detected when co-inoculated with either A-IN or A-ES, suggesting a reduced replication capacity in tomato plants if compared to the B-IN component (Figure 2). To determine whether the replication of the A component requires the presence of the B component, we performed local infections on tomato leaves using either the A-ES or the A-IN, individually. As shown in Figure S3, both A components were able to replicate autonomously; however, the replication efficiency of A-ES was consistently lower than that of A-IN.

**Figure 2.**
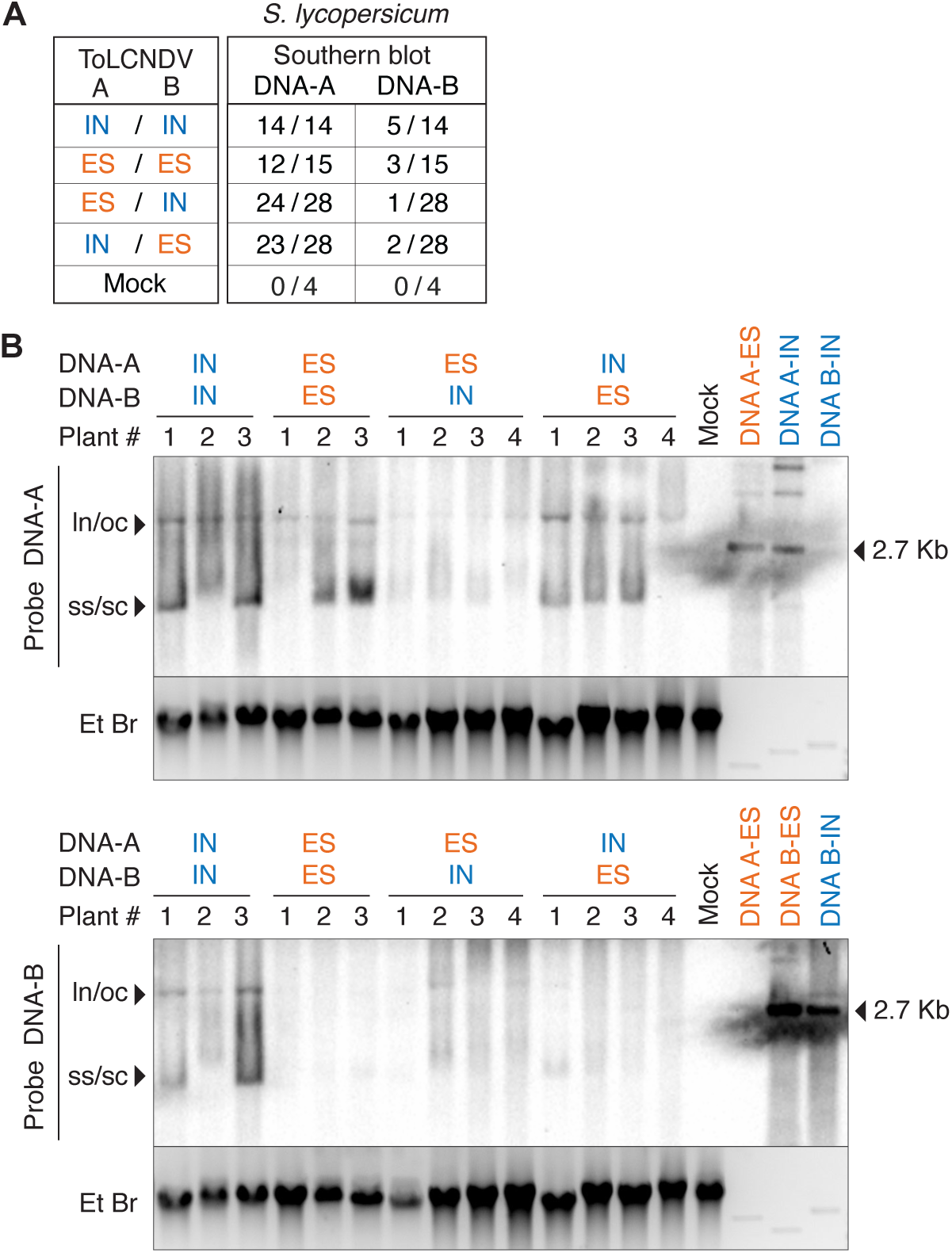
Detection of the India (IN) and Spanish (ES) ToLCNDV isolates and their pseudo-recombinants in local infection from tomato. (A) Summary of viral detection in agroinoculated tomato leaves at 5 dpi by Southern blot using DNA-A and DNA-B specific probes that recognize both the IN and ES ToLCNDV isolates. The table shows the number of plants testing positive for viral DNA relative to the total number of inoculated plants across three biological replicates. For each replicate, sample sizes ranged from 2 to 7 plants for the ToLCNDV-IN or ToLCNDV-ES, 8 to 12 plants for each pseudo-recombinant, and 1 to 2 plants for the mock control (binary vector lacking viral sequences). (B) Southern blot detection of viral DNA in agroinfiltrated tomato leaves at 5 days dpi. Approximately 1 µg of total genomic DNA per sample was separated on 0.8% agarose gels containing ethidium bromide, transferred to nylon membranes, and hybridized with DNA-A or DNA-B specific probes that detect both the IN and ES ToLCNDV isolates. The positions of the single-stranded (ss), supercoiled (sc), open circular (oc), and linear (ln) viral DNA forms, as well as the molecular-weight (M.W.) marker, are indicated. As an additional size reference, *Xba*I digested fragments from DNA-A-IN or DNA-B-ES infectious clones that released the corresponding monomeric viral genomic component (∼2.7 kb) were included on the right side of each gel. Samples shown correspond to one representative biological replicate.

To rule out the possibility that mutations in either genomic component were responsible for the ability of the pseudo-recombinant A-IN/B-ES to establish a systemic infection and induce symptoms in tomato (observed in 5 of the 14 plants that contained viral DNA), we performed Rolling Circle Amplification (RCA) on genomic DNA extracted from the five symptomatic plants, followed by nanopore sequencing (Figure S4A). Analysis of the A-IN component revealed multiple SNPs with a low proportion on the viral population; however, no consistent mutation was shared across all samples that could account for the infectivity of the A-IN/B-ES recombinant. Regarding the B-ES component, a full-length circular genome was identified in two of the five samples, while in a third sample, a defective DNA-B-ES molecule of approximately 1.5 kb was detected, containing a complete intergenic region (IR) and partial fragments of the BC1 and BV1 ORFs (Figure S4B). The absence of detectable B-ES in several of the plants infected with A-IN/B-ES, as shown by Southern blot and confirmed by RCA and sequencing, suggests two possible scenarios: either only a very limited number of B-ES molecules are able to reach the apical tissues, remaining below the detection threshold, or alternatively, A-IN is capable of systemically infecting tomato independently of the B component.

### Inefficient Systemic Spread of ToLCNDV-IN DNA-A in Tomato Highlights the Requirement for DNA-B

The low accumulation of the B component in A-IN/B-ES infected tomato plants, as revealed by Southern blot and RCA nanopore analyses, suggested that B-ES either fails to reach the apical tissues in detectable amounts or that A-IN is capable of systemic infection in tomato in the absence of DNA-B. To test whether the A component from the ToLCNDV-IN isolate can autonomously establish systemic infection in tomato, plants were agroinfiltrated with both genomic components or with DNA-A alone, and viral presence in apical tissues was assessed by molecular hybridization. No symptoms were observed, and squash blot analysis failed to detect viral DNA in plants inoculated solely with DNA-A (Figure S2). However, A-IN replicated efficiently in tomato leaves when inoculated alone (local infection; Figure S3), indicating that DNA-B is required for systemic infection. These observations were further examined using PCR, a more sensitive detection method than molecular hybridization, in plants agroinfiltrated with A-IN/B-IN, DNA-A alone, or the pseudo-recombinant A-IN/B-ES, which had previously been shown to infect tomato (Figures 1B and 1C). Viral DNA was detected by PCR in only 3 of 36 plants inoculated with DNA-A alone (Figure 3A), and in two of these cases the signal was faint and could not be confirmed by RCA nanopore sequencing (data not shown). On the other hand, DNA-A was readily detected by PCR in all plants agroinfiltrated with A-IN/B-ES, confirming that B-ES can provide the movement functions in trans required for systemic infection and that DNA-A from the IN isolate depends on the presence of DNA-B to successfully establish systemic infection in tomato (Figure 3B). These results indicated that DNA-A from the IN isolate is either incapable of systemic infection in tomato on its own or does so at very low efficiency (Figure 3A).

**Figure 3.**
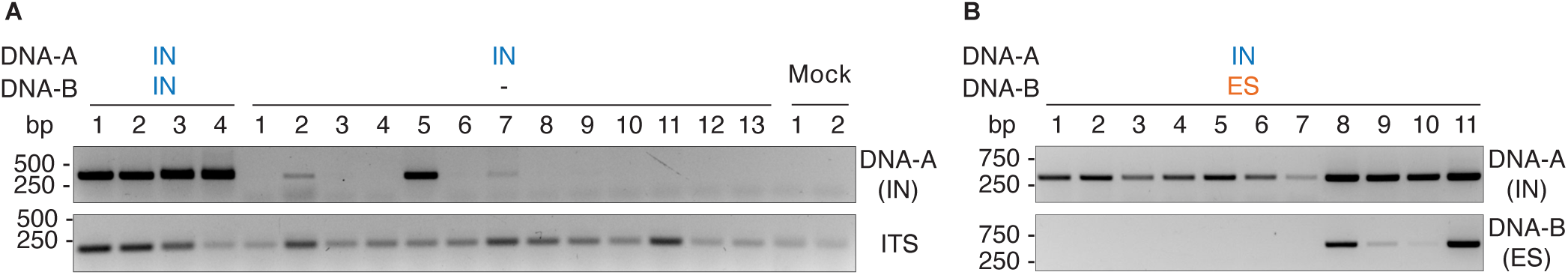
Detection of ToLCNDV-IN DNA-A in tomato when inoculated alone or with the heterologous DNA-B-ES. (A) Agarose gel electrophoresis showing PCR amplification of a fragment of the DNA-A-IN component from apical tomato leaves at 21 dpi when inoculated with its cognate DNA-A or by itself (upper panel). The Internal Transcribed Spacer (ITS) was amplified as an endogenous control. Detection was performed using a primer pair specific for the IN DNA-A component (lower panel). The gel displays representative samples from two independent experiments, which included a total of 36 plants inoculated with DNA-A-IN alone, 5 plants inoculated with the wild-type IN ToLCNDV isolate, 11 plants inoculated with the pseudo-recombinant A-IN/B-ES (see section B) and 6 plants for mock control (binary vector lacking viral sequences). Numbers indicate independent plants. (B) Agarose gel electrophoresis showing PCR amplification of fragments corresponding to the DNA-A-IN and DNA-B-ES components from apical tomato leaves at 21 dpi. Component-specific primer pairs were used to independently detect each genomic component. The gel displays the 11 samples agroinoculated with the pseudo-recombinant A-IN/B-ES. Numbers indicate independent plants.

### Pseudo-recombinant analysis of ToLCNDV-IN and ToLCNDV-ES in Nicotiana benthamiana plants, reveals complex interactions between their DNA components

In order to better understand the interplay between DNA-A and DNA-B from both isolates, we performed the same analysis with ToLCNDV pseudo-recombinants in the model host *Nicotiana benthamiana*. Plants were agroinfiltrated with either the infectious clones derived from the IN and ES isolates, or with the two potential pseudo-recombinants, A-IN/B-ES and A-ES/B-IN. Symptom development was monitored across two independent biological replicates (R1 to R2) with the four viral combinations and all the agroinfiltrated plants exhibited symptoms of varying severity (Table S1 and Figure S1). Overall, plants infected with constructs containing the A-IN component whether paired with B-IN or B-ES consistently developed much more severe symptoms than those infected with constructs containing the A-ES component. In both combinations, A-IN induced strong dwarfism and pronounced leaf curling (severity levels 5 and 4.5; 20/20 plants), whereas A-ES, even when paired with B-IN, produced only mild symptoms (severity levels 1.5–2; 20/20 plants) (Table S1 and Figure S1). Therefore, the ability to induce symptoms in *N. benthamiana* was primarily linked to the DNA-A component. The presence of viral DNA for of both components was confirmed in all *N. benthamiana* plants by molecular hybridization of squash blots using specific probes for DNA-A and DNA-B. DNA-A from each isolate was detected at comparable levels, regardless of the associated DNA-B component, indicating that A-IN and A-ES replicate with similar efficiency in *N. benthamiana* (Figure S5A and S5B). In contrast, the accumulation of the B component in systemic (apical) tissue depended on the associated A component. Thus, B-IN accumulated at higher level in the apical tissue when co-inoculated with its cognate A-IN than with A-ES, suggesting that the naturally occurring DNA-A/DNA-B pairing from the IN isolate was better adapted to *N. benthamiana* than the pseudo-recombinant combination (Figure S5B). Conversely, B-ES accumulated to similar levels whether associated with any of the A components (Figure S5B). We previously demonstrated that DNA-A from the IN isolate is either incapable of systemically infecting tomato on its own or does so with very low efficiency (Figure 3). To determine whether this limitation is host-specific, we evaluated its infectivity in *N. benthamiana*. Plants were agroinfiltrated with the DNA-A component from either the IN or ES isolates, either alone or in combination with their cognate DNA-B component. Symptom development and viral accumulation in apical tissues was monitored across two independent biological replicates. Plants inoculated with A-ES alone remained asymptomatic, while those infected with A-IN exhibited very mild symptoms (average severity score: 1.5 out of 5; Table S1). Viral DNA accumulation from the A components was assessed by squash blot hybridization of apical leaf petioles. Viral DNA was detectable in all samples (Figure S5B). In the case of A-ES, signal intensity was comparable to that observed in plants inoculated with the DNA-A and DNA-B combination. However, in A-IN-inoculated plants, the hybridization signal was notably weaker than in plants receiving both A and B components of the IN isolate, suggesting lower levels of viral accumulation. Given the semi-quantitative nature of squash blotting, we performed another independent biological replicate and performed qPCR analysis to obtain more accurate measurements of viral load. Plants agroinfiltrated with either DNA-A alone or with its cognate DNA-B as a control, were analysed. As shown in Figure S6, viral accumulation in plants infected with the naturally occurring IN isolate was significantly higher than in those inoculated with A-IN alone (Figure S6A). A similar trend was observed for the ES isolate, indicating that both isolates infect *N. benthamiana* more efficiently when co-inoculated with their cognate DNA-B component (Figure S6B).

These results indicated that the reduced symptom severity observed in A-IN-infected N. benthamiana is likely attributable to both the absence of the DNA-B component and the overall lower levels of viral accumulation. Thus, while the IN isolate can establish a systemic infection in N. benthamiana as a monopartite virus, it did so with high efficiency (100% of the plants were infected) but with a drastic reduction of the viral titre and symptom severity. Likewise, the ES isolate also displayed a monopartite-like infection pattern in *N. benthamiana*, though without inducing any visible symptoms. Together, these findings suggested that both ToLCNDV-IN and ToLCNDV-ES are capable of replicating and moving systemically in N. benthamiana without their cognate DNA-B components, although less effectively than when both components are present.

### Systemic Infection by ToLCNDV-ES in *N. benthamiana* required the Coat Protein

It has been previously shown that bipartite begomoviruses with mutations or replacements in the CP can still establish systemic infections, although typically associated with mild symptoms, since the proteins responsible for viral movement are encoded in the DNA-B component (Brough et al. 1988; Pooma et al. 1996; Sudarshana et al. 1998). Although mutations in the CP from ToLCNDV-IN, reduce ssDNA accumulation, the capside protein is not required for systemic infection and symptom development (Padidam et al. 1995, 1996). To generate an infectious ToLCNDV-ES-based construct capable of serving as a reporter for viral cell-to-cell and systemic movement, we engineered a modified version of the DNA-A component (DNA-A^ΔCP^) in which the DsRed gene was inserted into a 1.2 mer clone of DNA-A from ToLCNDV-ES (Figure 4A). To avoid disrupting the overlapping AV2 gene, we replaced the last 591 bp of the AV1 gene with the DsRed ORF, resulting in an in-frame fusion between DsRed and the N-terminal 60 amino acids of the CP. This construct should allow the expression of DsRed from the native AV2/AV1 promoter located in the intergenic region (IR) whereas will generate a DNA-A clone deleted for CP (DNA-A^ΔCP^).

**Figure 4.**
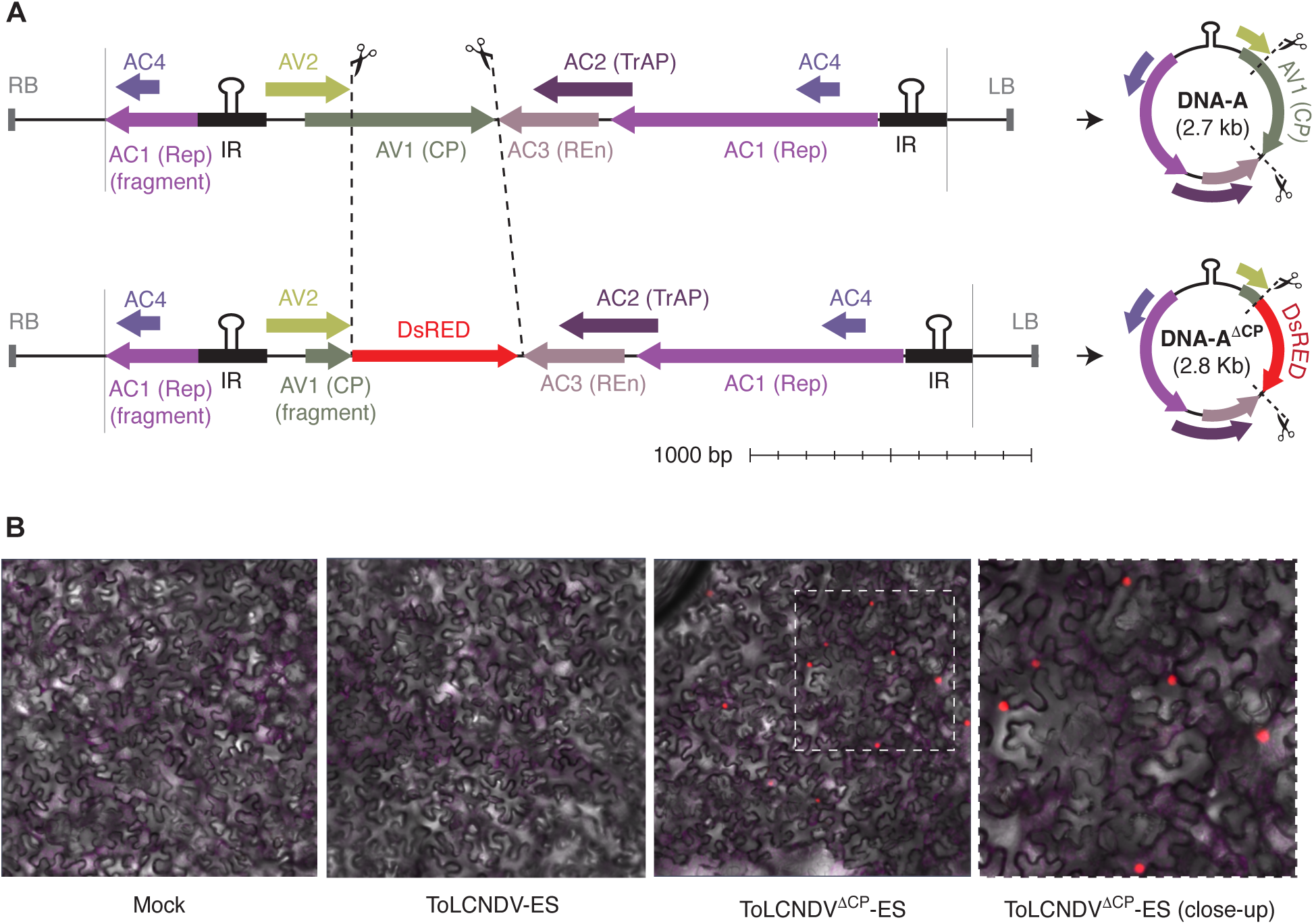
Detection of DsRed-tagged ToLCNDV DNA-A clone in *N. benthamiana* leaves. (A) Schematic illustration of the modified DNA-A-ES component (DNA-A^ΔCP^), in which the last 483-nt of the CP gene were replaced by the DsRed gene in a 1.2-mer infectious clone of ToLCNDV-ES. Linear and circular visualization of the wt and recombinant DNA-A^ΔCP^-ES components are shown. (B) Confocal laser scanning microscope (CLSM) images of epidermal cells of N. benthamiana leaves agroinoculated with ToLCNDV-ES or the recombinant clone ToLCNDV^ΔCP^-ES at 3 days post-agroinfiltration (dpa).

To confirm if DsRed was expressed, we agroinfiltrated *N. benthamiana* leaves with either the wild-type virus or the DNA-A^ΔCP^ construct with its cognate DNA-B and monitored fluorescence at 3 days post-infiltration (dpi) in the infiltrated leaves and at 21 dpi in the apical tissues.

Notably, DsRed acts as a cell-autonomous reporter because it forms obligate tetramers, producing a complex too large to pass through plasmodesmata (Nishizawa et al. 2006; Wall et al. 2000). Consequently, the appearance of DsRed fluorescence in apical would serve as a clear indicator that the virus has successfully established a systemic infection. At 3 dpi, DsRed fluorescence was detected in the nuclei of multiple cells in the infiltrated area, confirming the expression of the fluorescent protein from the viral promoter (Figure 4B). Surprisingly, no DsRed signal was observed in the DNA-A^ΔCP^/DNA-B-infected apical leaves (data not shown), suggesting that this CP-mutated version of ToLCNDV-ES was not able to systemically infect the plant.

To investigate whether this lack of signal was due to a failure in systemic movement of the ΔCP viral clone, we analysed viral DNA accumulation in apical tissues at 21 dpi in two independent biological replicates. While wild-type ToLCNDV-ES, or even the DNA-A component alone, could be detected systemically (previously shown, Figure S6), no viral DNA was identified in plants agroinfiltrated with DNA-AΔCP, regardless of whether it was co-infiltrated alone or with its cognate DNA-B component (Figure 5A). These results suggested that, at least in *N. benthamiana*, ToLCNDV-ES did not behave as a canonical bipartite begomovirus, and that the CP was still essential for viral movement and systemic infection. Moreover, the movement proteins encoded by DNA-B seem to increase, rather than substitute for CP function, as viral accumulation was significantly reduced in the absence of DNA-B as obtained by qPCR (Figure 5B and S6).

**Figure 5.**
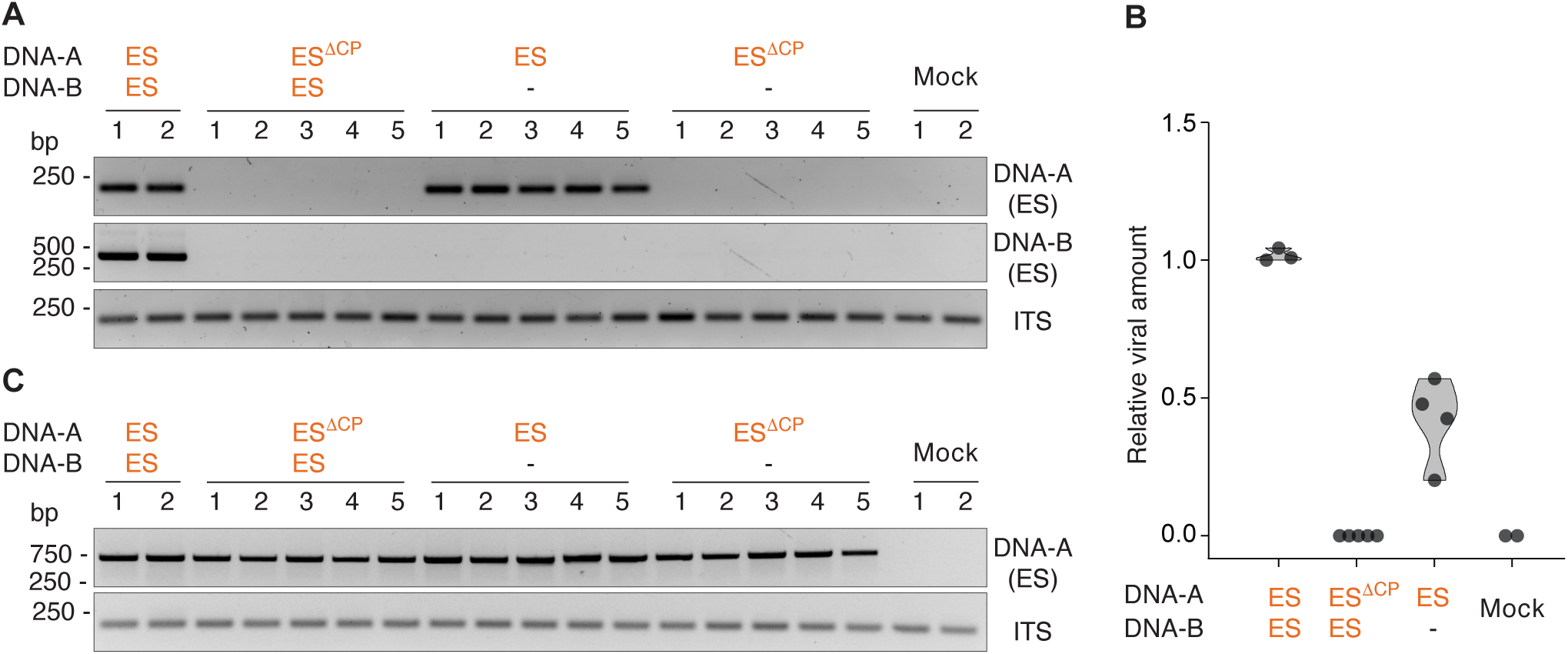
Detection of ToLCNDV-ES DNA-A and DNA-A^ΔCP^ in *N. benthamiana* when inoculated with or without its cognate DNA-B component. (A) Agarose gel electrophoresis showing PCR amplification of fragments corresponding to the DNA-A and DNA-B components from apical *N. benthamiana* leaves at 21 days post-inoculation (dpi). Component-specific primers were used for each PCR. The Internal Transcribed Spacer (ITS) was used as an endogenous control. The gel shows representative samples from one biological replicate out of two independent replicates. (B) Relative viral accumulation in systemic tissue (21 dpi) of *N. benthamiana* plants measured by qPCR with primers for the DNA-A-ES component. The graph displays the results from one biological replicate, with each dot representing an individual plant. (C) Agarose gel electrophoresis showing PCR amplification of fragments corresponding to ToLCNDV-ES DNA-A using episomal primers in agroinfiltrated *N. benthamiana* leaves collected at 5 dpi. Numbers indicate independent plants. The mock samples come from plants inoculated with binary vector lacking viral sequences

To rule out the possibility that CP deletion affected viral replication, we assessed local viral accumulation in agroinfiltrated leaves using PCR with episomal-specific primers for DNA-A. The ΔCP viral mutant accumulated to similar levels as the wild-type virus, indicating that the lack of systemic movement in the absence of CP, even when DNA-B-encoded movement proteins were present, was not due to impaired replication (Figure 5C).

## Acknowledgements

We kindly thank Supriya Chakraborty (Jawaharlal Nehru University, India) for providing the agroinfective clones for ToLCNDV-IN. This work was funded by Plan Nacional I + D + i, Ministerio de Economía y Competitividad (Agencia Estatal de Investigación), Spain (PID2019-107657RB-C22; PID2022-139376OB-C31). The funders had no role in study design, data collection and analysis, decision to publish, or preparation of the manuscript.

---

**Table S1.** Symptoms development in tomato and *Nicotiana benthamiana* plants inoculated with individual or combined genomic components of the two ToLCNDV isolates (IN and ES) and their pseudo-recombinants. The table shows the number of symptomatic plants relative to the total number of plants inoculated from five independent replicates (R1 to R5) at 21-35 dpi. The number of symptomatic plants for each combination are indicated between brackets. Symptom severity was scored on a scale from 1 to 5, where * indicates mild symptoms (1.5–2) and ** indicates very severe symptoms (4–5).

|  | ToLCNDV | Symptoms |  |  |  |  |  |
| --- | --- | --- | --- | --- | --- | --- | --- |
|  | A / B | R1 | R2 | R3 | R4 | R5 | Total |
| **Tomato** | IN / IN | 10/10 (2) | 10/10 (4) | 5/5 (5) | 10/10 (5) | 10/10 (5) | 45/45** |
|  | ES / ES | 0/10 | 0/10 | 0/5 | 0/10 | 0/17 | 0/52 |
|  | ES / IN | 0/10 | 0/10 | 0/8 | 0/10 | 0/20 | 0/58 |
|  | IN / ES | 3/10 (4) | 0/10 | 1/8 (5) | 1/10 (5) | 0/20 | 5/58** |
|  | IN / - | - | 0/5 | - | - | - | 0/5 |
|  | ES / - | - | 0/5 | - | - | - | 0/5 |
|  | Mock | 0/4 | 0/5 | 0/3 | 0/10 | 0/10 | 0/32 |
| ***N.benthamiana*** | IN / IN | 10/10 (5) | 10/10 (5) | 3/3 (5) | 3/3 (5) | 10/10 (5) | 36/36** |
|  | ES / ES | 10/10 (1.5) | 10/10 (1.5) | 3/3 (1.5) | 3/3 (2.5) | 10/10 (1.5) | 36/36* |
|  | ES / IN | 10/10 (2) | 10/10 (2) | 3/3 (1.5) | 3/3 (1.5) | 10/10 (2) | 36/36* |
|  | IN / ES | 10/10 (4.5) | 10/10 (4.5) | 3/3 (5) | 3/3 (5) | 10/10 (4.5) | 36/36** |
|  | IN / - | - | 5/5 (1.5) | - | - | - | 5/5* |
|  | ES / - | - | 0/5 | - | - | - | 0/5 |
|  | Mock | 0/7 | 0/5 | 0/3 | 0/2 | 0/8 | 0/25 |

**Figure S1.**
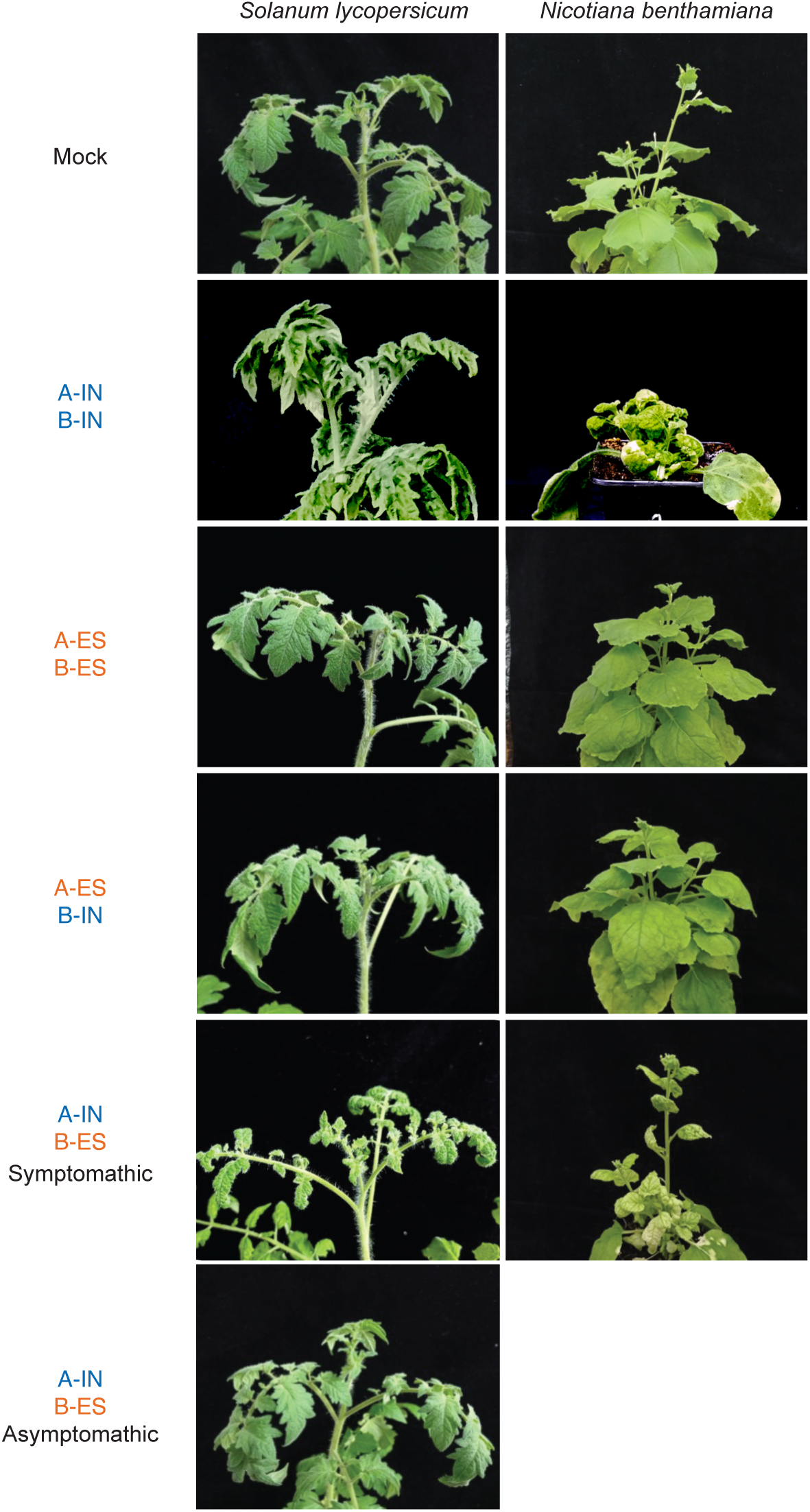
Symptom development in tomato and *N. benthamiana* plants inoculated with ToLCNDV isolates IN and ES and their pseudo-recombinants. Representative symptom development in tomato and N. benthamiana plants inoculated with both IN and ES ToLCNDV isolates and their corresponding pseudo-recombinants (A-IN/B-ES and A-ES/B-IN) at 24 dpi. For the pseudo-recombinant A-IN/B-ES a symptomatic and an asymptomatic representative plant are shown. Mock plants were inoculated with binary vector lacking viral sequences

**Figure S2.**
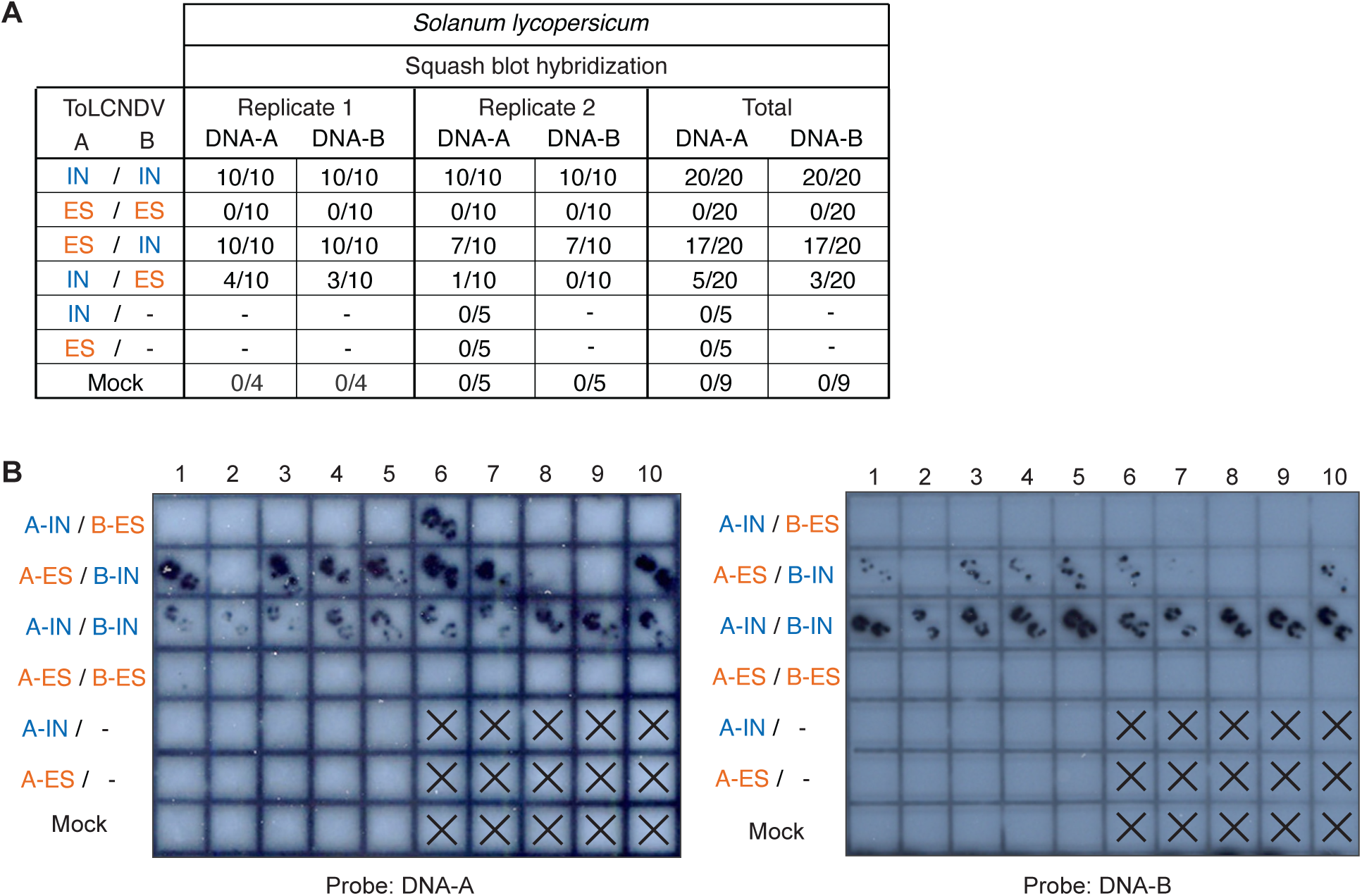
Detection of ToLCNDV-IN, ToLCNDV-ES, and their pseudo-recombinants in tomato tissues by squash blot hybridization. (A) Summary of viral detection by squash blot hybridization from apical tomato leaves at 24 dpi. The table shows the number of plants testing positive for viral DNA relative to the total number of inoculated plants across two biological replicates (R1 and R2). (B) Squash blot detection of viral DNA in apical tomato leaves at 24 dpi. Squash blots were prepared from cross-sections of petioles from young systemic leaves of tomato plants performed on positively charged nylon membranes. One representative replicate is shown. Blots were hybridized with DNA-A and DNA-B specific probes that recognize both the IN and ES ToLCNDV isolates. Mock plants were inoculated with a binary vector lacking viral sequences.

**Figure S3.**
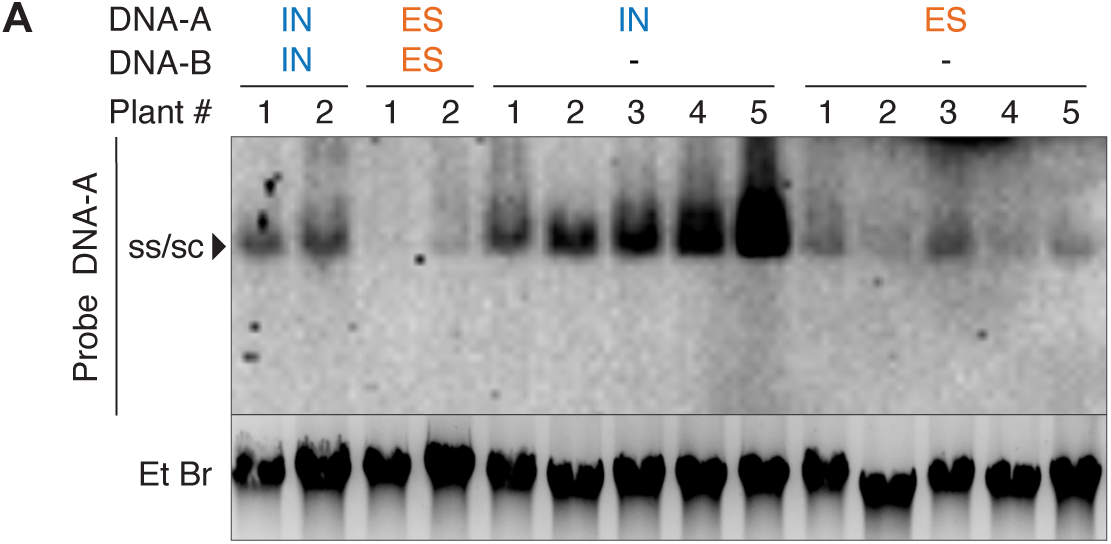
Detection of ToLCNDV IN and ES isolates and their pseudo-recombinants in inoculated tomato leaves by Southern blot. Southern blot detection of viral DNA in infiltrated tomato leaves at 5 dpi. Approximately 1 µg of total genomic DNA per sample was separated on 0.8% agarose gels containing ethidium bromide, transferred to nylon membranes, and hybridized with DNA-A specific probe that detect DNA-A from both the IN and ES ToLCNDV isolates. The position of the single-stranded (ss) and supercoiled (sc) DNA forms are indicated. Numbers indicate independent plants. The blot shows one representative replicate from two independent experiments.

**Figure S4.**
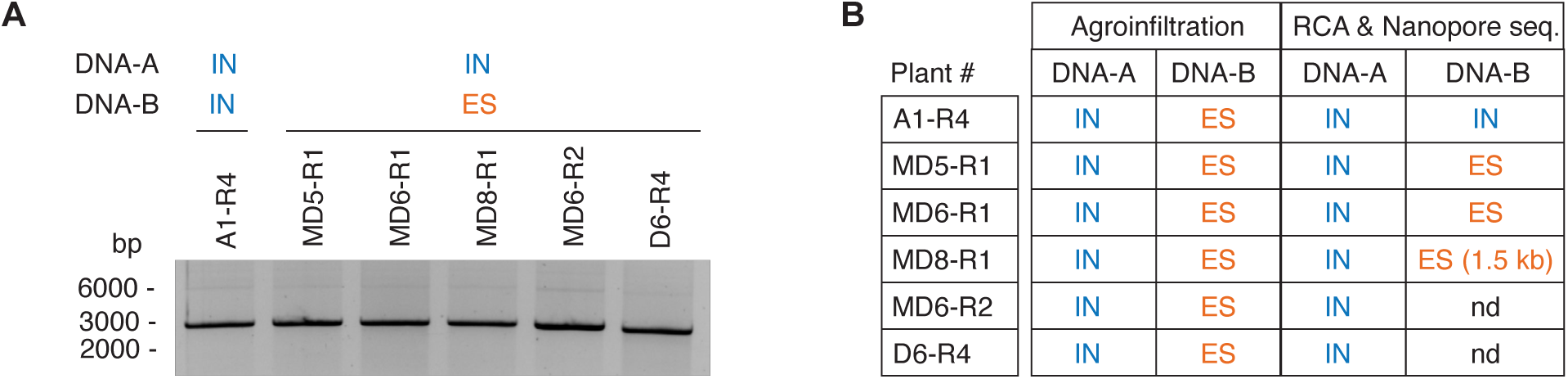
Rolling circle amplification (RCA) amplification and nanopore sequencing of genomic components from symptomatic tomato plants infected with the pseudo-recombinant A-IN/B-ES. (A) RCA from five tomato plants inoculated with A-IN/B-ES and sampled at 21 dpi. RCA products were digested with *Xba*I and resolved by electrophoresis on a 1% agarose gel, yielding the expected linear fragments corresponding to DNA-A and/or DNA-B. A sample from a tomato plant infected with ToLCNDV-IN isolate (A1-R4) was included as a positive control for restriction digestion and banding pattern reference. (B) Summary for the genomic component identified after RCA and nanopre sequencing techomology for each plant inoculated with the indicated viralgenome combination. “nd” indicates that the DNA-B component was non-detected on that sample.

**Figure S5.**
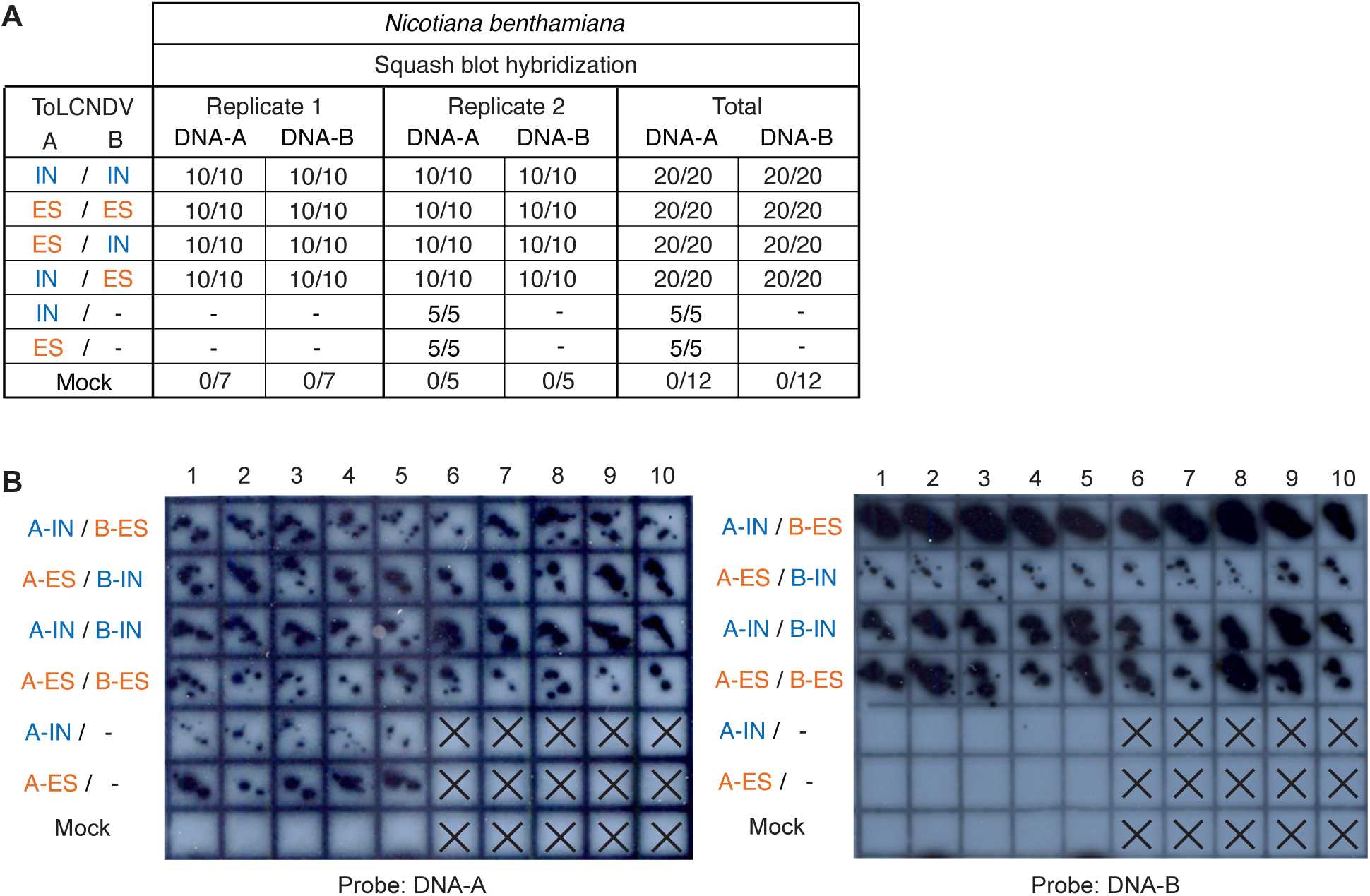
Detection of ToLCNDV-IN, ToLCNDV-ES, and their pseudo-recombinants in *N. benthamiana* tissues by squash blot hybridization. (A) Summary of viral detection by squash blot hybridization from apical *N. benthamiana* leaves at 24 dpi. The table shows the number of plants testing positive for viral DNA relative to the total number of inoculated plants across two biological replicates (R1 and R2). (B) Squash blot detection of viral DNA in apical *N. benthamiana* leaves at 24 dpi. Squash blots were prepared from cross-sections of petioles from young systemic leaves of *N. benthamiana* plants performed on positively charged nylon membranes. One representative replicate is shown. Blots were hybridized with DNA-A and DNA-B specific probes that recognize both the IN and ES ToLCNDV isolates. Mock plants were inoculated with a binary vector lacking viral sequences.

**Figure S6.**
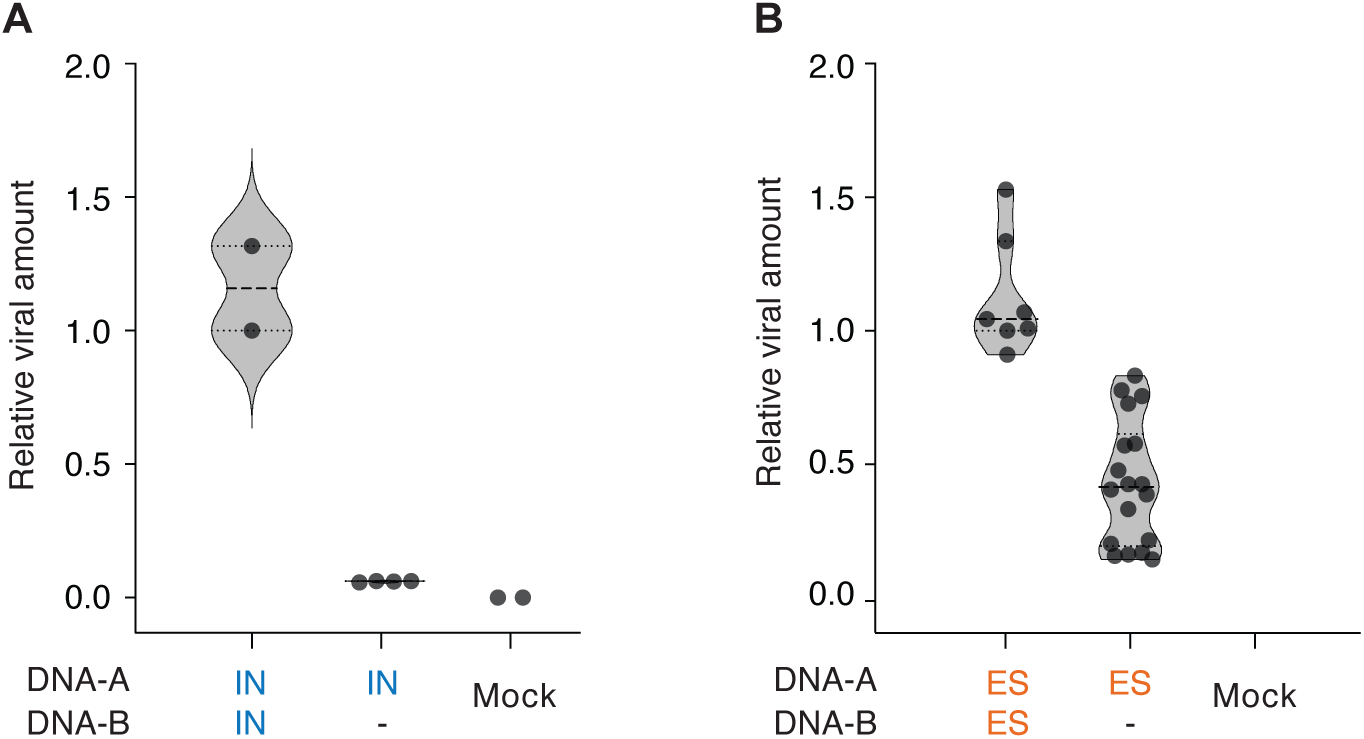
Detection of ToLCNDV DNA-A in N. benthamiana when inoculated with or without its cognate DNA-B component. Relative viral accumulation from systemic tissue (21 dpi) of N. benthamiana plants measured by qPCR with primers specific for DNA-A-IN (A) and for DNA-A-ES (B). The graph displays the results from one biological replicate in (A) or three biological replicates in (B), with each dot representing an individual plant. The mock samples come from plants inoculated with binary vector lacking viral sequences.

